# A Deep Learning Framework for Prediction of Clinical Drug Response of Cancer Patients and Identification of Drug Sensitivity Biomarkers using Preclinical Samples

**DOI:** 10.1101/2021.07.06.451273

**Authors:** David Earl Hostallero, Lixuan Wei, Liewei Wang, Junmei Cairns, Amin Emad

## Abstract

**Background:** Prediction of the response of cancer patients to different treatments and identification of biomarkers of drug sensitivity are two major goals of individualized medicine. In this study, we developed a deep learning framework called TINDL, completely trained on preclinical cancer cell lines, to predict the response of cancer patients to different treatments. TINDL utilizes a tissue-informed normalization to account for the tissue and cancer type of the tumours and to reduce the statistical discrepancies between cell lines and patient tumours. In addition, this model identifies a small set of genes whose mRNA expression are predictive of drug response in the trained model, enabling identification of biomarkers of drug sensitivity.

**Results:** Using data from two large databases of cancer cell lines and cancer tumours, we showed that this model can distinguish between sensitive and resistant tumours for 10 (out of 14) drugs, outperforming various other machine learning models. In addition, our siRNA knockdown experiments on 10 genes identified by this model for one of the drugs (tamoxifen) confirmed that all of these genes significantly influence the drug sensitivity of the MCF7 cell line to this drug. In addition, genes implicated for multiple drugs pointed to shared mechanism of action among drugs and suggested several important signaling pathways.

**Conclusions:** In summary, this study provides a powerful deep learning framework for prediction of drug response and for identification of biomarkers of drug sensitivity in cancer.

## INTRODUCTION

Cancer is one of the deadliest public health problems worldwide and cases are still rapidly growing. In 2020, it is estimated that around 10 million people have died of cancer [1]. Individualized medicine is a promising concept, which aims to improve the prognosis of patients by adapting the patient’s treatment to their unique clinical and molecular characteristics. One of the main goals of individualized medicine is the prediction of the patients’ response to different treatments, and identification of biomarkers that enable such predictions. High throughput sequencing technologies along with major initiatives such as The Cancer Genome Atlas (TCGA) [2] have provided a unique opportunity for machine learning (ML) algorithms to address these challenges. However, ML models and particularly deep learning (DL) approaches require a large number of samples with known drug response to train generalizable models. However, data on clinical drug response (CDR) of cancer patients, even in large databases such as TCGA, is usually small for the majority of the drugs and does not lend itself to training of DL models.

On the other hand, large databases of molecular profiles of hundreds of *in-vitro* cancer cell lines (CCLs) and their response to hundreds of drugs [3–5] have enabled development of various ML algorithms for prediction of drug response [6–8]. Unfortunately, these models, even though accurate in predicting the drug response of held-out CCLs, usually do not generalize well to predicting the CDR of real tumours from cancer patients, and their prediction performance significantly deteriorates due to the major biological and statistical differences between CCLs and tumours [9].

Recognizing these issues, some studies have adopted to utilize tumour samples with known CDR in the training of their models, either by fully training their models on data corresponding to tumour samples [10–12], or by using them in addition to CCLs (e.g., using transfer learning [13]). However, as a result of this strategy, these studies have only been able to develop models on very few drugs due to the small samples sizes of patient cohort data with known drug response. Another strategy is to train ML models completely on preclinical CCLs but use computational approaches to overcome the statistical differences between CCLs and tumours. For example, multiple approaches [9, 14] have used batch removal methods such as ComBat [15] to reduce the discrepancy between the training CCLs and test tumours. One limitation of these methods is that ComBat is used as a preprocessing step such that the gene expression profile of both CCLs (training set) and tumours (test set) are adjusted. As a result, prediction of CDR of new cancer patients requires re-training of the model.

In this study, our goal was to develop a deep learning computational pipeline, fully trained on gene expression profile and drug response of preclinical CCLs, to 1) predict the CDR of cancer patients and 2) to identify biomarkers of drug sensitivity for a variety of cancer drugs. Motivated by Huang et al. [9], who showed that carefully incorporating information on the tissue (or cancer) type of the test samples can improve the predictive power of computational models, we developed a <u>d</u>eep learning pipeline with <u>t</u>issue-informed <u>n</u>ormalization (TINDL) to achieve these goals. Unlike methods mentioned above, TINDL requires normalization of only test samples, and as a result re-training of the model is not necessary for new test samples.

The TINDL pipeline includes two phases. The first phase is responsible for prediction of CDR of cancer patients, while the second phase utilizes these predictions to identify a small number of genes that considerably contribute to the predictive ability of the model. Focusing on drugs shared between the Genomics of Drug Sensitivity in Cancer (GDSC) [3] and TCGA [2], we showed that TINDL can distinguish between the sensitive and resistant patients for 10 (out of 14) drugs, considerably improving the performance of other methods, including our previous work TG-LASSO [9]. TINDL utilizes a simple, yet effective, tissue-informed normalization to reduce the statistical discrepancies between the gene expression profile of the training and test samples. We showed that TINDL outperforms other DL-based models that try to explicitly remove these discrepancies using other techniques such as ComBat or domain adaptation [16, 17].

Focusing on tamoxifen, for which TINDL performed best, we showed that only a small panel of genes identified by TINDL can be used to predict the CDR of cancer patients. Moreover, using siRNA gene knockdown of 10 genes identified by TINDL in a breast cancer cell lines (MCF7), we showed that the knockdown of any of these genes significantly changes the response to tamoxifen. These *in-vitro* experiments further validate the TINDL pipeline and its ability to identify biomarkers of drug sensitivity.

## RESULTS

### Deep learning prediction of clinical drug response of cancer patients and identification of biomarkers of drug sensitivity using *in vitro* cell line data

We developed a <u>d</u>eep learning pipeline with <u>t</u>issue-informed <u>n</u>ormalization (called TINDL) to 1) predict the clinical drug response (CDR) of cancer patients (test set) and 2) identify predictive biomarkers of drug sensitivity based on models completely trained on preclinical cell line data (training set). The pipeline has two major phases: the modeling phase and the gene identification phase. In the modeling phase (Figure 1a), a neural network is trained using the gene expression (GEx) profiles of cancer cell lines (CCLs) and their response to a drug (e.g., normalized ln(IC50) values in this study). The trained model was then used to predict the drug response of cancer patients based on the carefully normalized GEx profiles of their primary tumours. Details of the DL architecture are provided in Methods.

**Figure 1.**
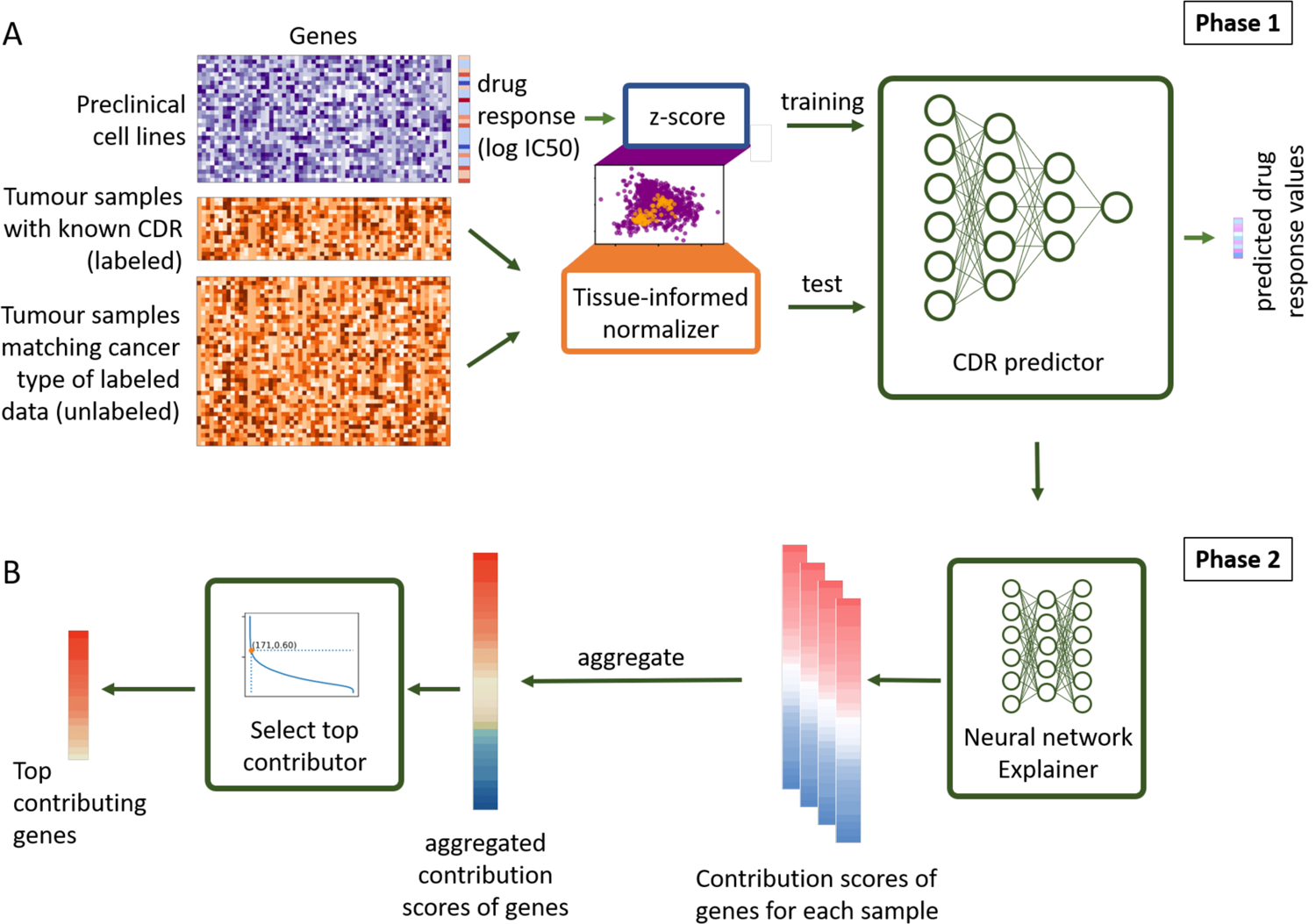
The pipeline used for prediction of drug responses and identification of important genes. In Phase 1, (A) the gene expression data of the cancer cell lines (CCL) and log IC50 are both z-score normalized, while the tumour gene expression (test data) is normalized using the tissue-informed normalizer. We then used the train a cancer drug response (CDR) predictor using the CCL data. After training, we predict the response value for the tumours. In Phase 2 (B), we take the trained CDR predictor and train a neural network explainer using the same training data. We used the trained explainer to give gene contribution scores for each genes of the test samples. We aggregated the scores across samples and then selected the top genes by estimating the point of maximum curvature.

We designed the normalization step of GEx profiles of patient tumours to address two important issues. First, we required this approach to remove the discrepancy between the statistical properties of GEx of CCLs and patient tumours, originating from the technical differences in protocols for measuring the data and the biological differences between pre-clinical CCLs and clinical tumours. Second, we required this approach to incorporate information on the tissue of origin (or cancer type) of tumours in the prediction task. In a previous study [9], we showed that information on the tissue of origin of samples plays an important role in improving prediction performance, however most commonly used methods for this task are not capable of appropriately incorporating this information. For this purpose and given a drug, we first identified the set of tissues (henceforth referred to as “target tissues”) of the clinical samples to whom the drug was administered. Then, we collected additional GEx profile of samples from the same target tissues, independent of what drug was used for their treatment. The GEx profile of each test sample was then normalized against this additional set of “unlabeled” data (see Methods for details).

This simple, yet effective, normalization approach used in our pipeline removes the statistical discrepancy between the test and training datasets by mapping the expression of each gene in each dataset to a distribution with unit variance and zero mean. However, since the test samples are normalized while considering the GEx of a much larger unlabeled set of samples, this normalization will not be negatively affected if the size of the test set is small (e.g., if we want to predict the drug response of a single sample). This is different from methods that perform the normalization using only the test samples. In addition, since the normalization is done independently for the training and test sets, one does not need to retrain the DL model every time the drug response of a new test sample is to be predicted (a shortcoming of our previous approach [9]).

The second phase of the pipeline seeks to assign a contribution score to each gene based on its contribution to the trained predictive model. In this phase, we first use CXPlain [18] to assign a sample-specific score to each gene. These scores are then averaged over all samples (separately for each gene) and normalized to provide a final contribution score. Additionally, we use the distribution of these scores to systematically identify the critical point that the contribution of the genes diminishes, enabling us to narrow down the top ranked list of genes for follow-up analysis (e.g., pathway enrichment analysis, gene knockdown experiments, etc.). The details of this phase are provided in Methods.

### TINDL can distinguish between sensitive and resistant patients for the majority of the drugs

In order to assess the performance of TINDL in predicting CDR of cancer patients, we obtained GEx profile of primary cancer tumours from the TCGA database [2]. We used data corresponding to RECIST CDR of TCGA patients carefully collected in [10] and identified 14 drugs that satisfy two conditions: 1) there were at least 20 patients with known CDR values for each drug in TCGA database and 2) the ln(IC50) response of these drugs were measured in the GDSC database. Similar to previous studies [9, 14], we transformed the CDR of these tumours into a Boolean label in which “resistant” referred to patients with CDR of “stable disease” or “progressive disease” and “sensitive” referred to patients with CDR of “complete response” or “partial response”. These CDR values were used to evaluate the predicted drug response values using TINDL and other algorithms but were not used for training them. The list of these 14 drugs, number of TCGA patients, and their cancer type are provided in Supplementary Table S1. Similarly, we obtained GEx profiles and ln(IC50) drug response of CCLs from different lineages from the GDSC database [3], corresponding to the 14 drugs above (See Supplementary Table S1 for number of training samples for each drug).

Following previous work in this area [9, 14], we used a one-sided Mann-Whitney U test to determine if the predicted ln(IC50) values of resistant patients for a drug are significantly higher than sensitive patients. Table 1 shows the performance of TINDL in prediction of CDR of TCGA samples using preclinical GDSC samples based on this statistical test for different drugs (also see Supplementary Table S2 for the area under the receiver operating characteristic (AUROC) values). TINDL is capable of distinguishing between resistant and sensitive patients for 10 (out of 14) drugs (p < 0.05, one-sided Mann-Whitney U test) with a combined p-value of 2.77 E-10 (Fisher’s method).

**Table 1:** The number of TCGA samples and the performance of TINDL in predicting their CDR for 14 drugs. The first column shows the name of the drug, the second column shows the total number of clinical samples for each drug, and third and fourth columns show the number of sensitive and resistant samples, respectively. The fifth column shows the p-value of a one-sided Mann-Whitney U test to determine if TINDL can distinguish between sensitive and resistant patients. To ensure the results are not biased by the initialization of the model’s parameters, TINDL was trained using ten random initializations and the mean aggregate of its prediction were used to calculate the p-values. Drugs are sorted based on their associated p-value.

| Drug | Number of clinical samples | Number of sensitive samples | Number of resistant samples | p-value |
| --- | --- | --- | --- | --- |
| Cisplatin | 303 | 237 | 66 | 6.36E-4 |
| Tamoxifen | 20 | 14 | 6 | 1.14E-3 |
| Etoposide | 84 | 73 | 11 | 4.00E-3 |
| Doxorubicin | 100 | 68 | 32 | 1.42E-2 |
| Paclitaxel | 158 | 111 | 47 | 2.29E-2 |
| Vinorelbine | 30 | 23 | 7 | 2.41E-2 |
| Oxaliplatin | 54 | 33 | 21 | 2.41E-2 |
| Temozolomide | 95 | 11 | 84 | 2.94E-2 |
| Bleomycin | 52 | 46 | 6 | 3.41E-2 |
| Gemcitabine | 157 | 75 | 82 | 4.57E-2 |
| Cyclophosphamide | 101 | 96 | 5 | 5.60E-2 |
| Pemetrexed | 38 | 18 | 20 | 2.86E-1 |
| Irinotecan | 23 | 6 | 17 | 3.04E-1 |
| Docetaxel | 102 | 67 | 35 | 7.04E-1 |

Next, we defined a measure called precision at k^th^ percentile to determine whether patients whose predicted ln(IC50) is within the lower tail of the distribution correspond to sensitive patients (i.e. responders to the drug). For different values of k, tumours with predicted ln(IC50) in the bottom k% were predicted as sensitive and their count was used to calculate precision. Figure 2A and Supplementary Table S3 show precision at k^th^ percentile of TINDL for different values of k. These results suggest that for six drugs (tamoxifen, etoposide, vinorelbine, cyclophosphamide, bleomycin, and cisplatin) TINDL can identify responders with a precision at kth percentile above 84% for any choice of k. The distribution of predicted CDR values for sensitive and resistant patients for these drugs are shown in Figure 2B.

**Figure 2:**
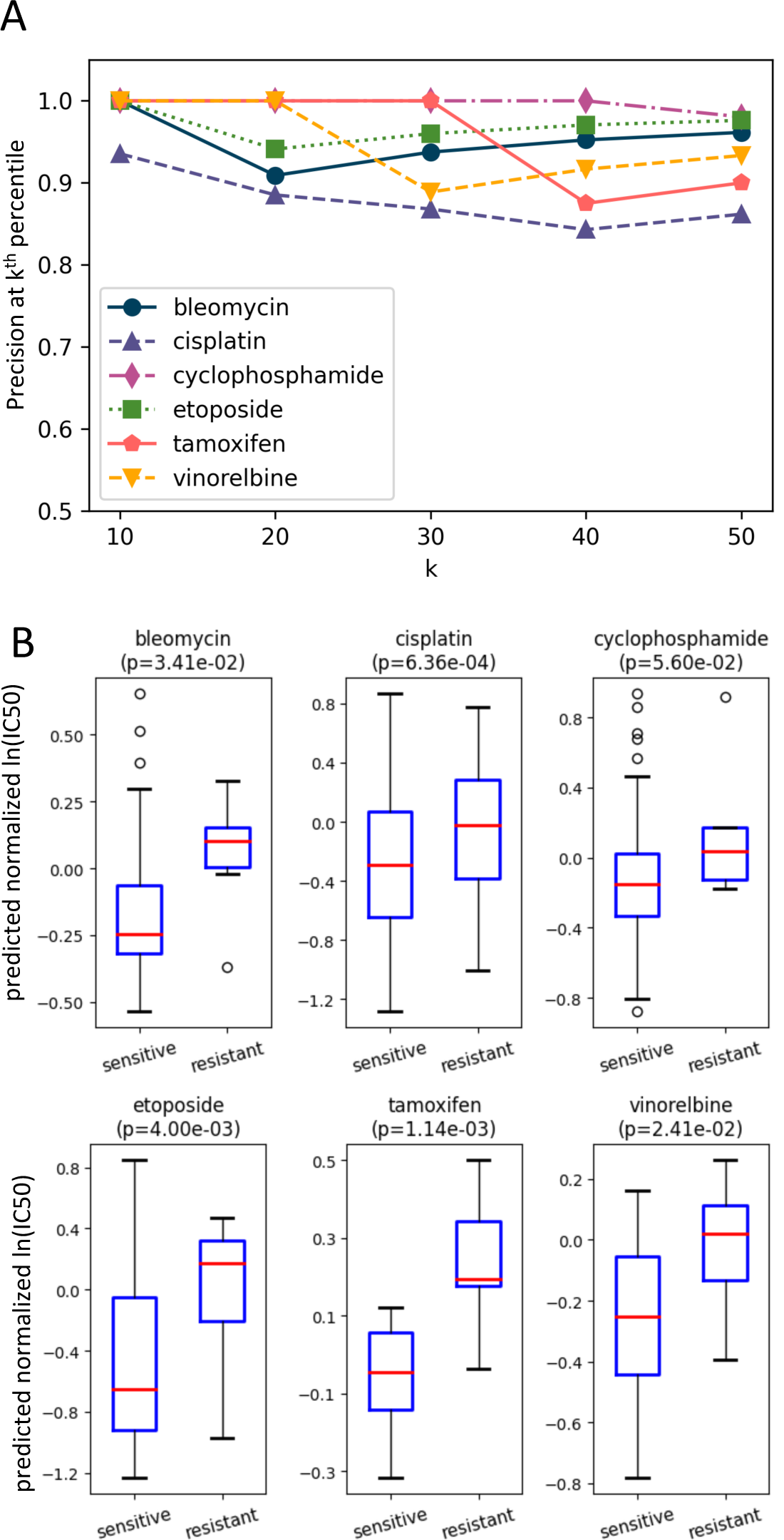
Performance metrics for a subset of the drugs. To prevent the figure from becoming cluttered, the results corresponding to only six drugs are shown (see Supplementary Tables S2 and S3 for performance metrics of all drugs). A) Precision at kth percentile for identification of sensitive patients. B) Distribution of predicted drug response for sensitive and resistant patients. The p-values are calculated using a one-sided Mann Whitney U test.

### TINDL outperforms alternative methods in prediction of clinical drug response

Next, we sought to determine how TINDL performs against alternative computational models. For this purpose, we considered multiple traditional and state-of-the-art machine learning models [9, 14] for prediction of CDR of cancer patients from preclinical CCLs. The detailed performance measures for each drug and each model are provided in Supplementary Table S2 and the summary of the results are provided in Table 2. In this table, we used the combined p-value of 14 drugs to summarize the performance of different methods (Fisher’s method for combining p-values was used).

**Table 2:**
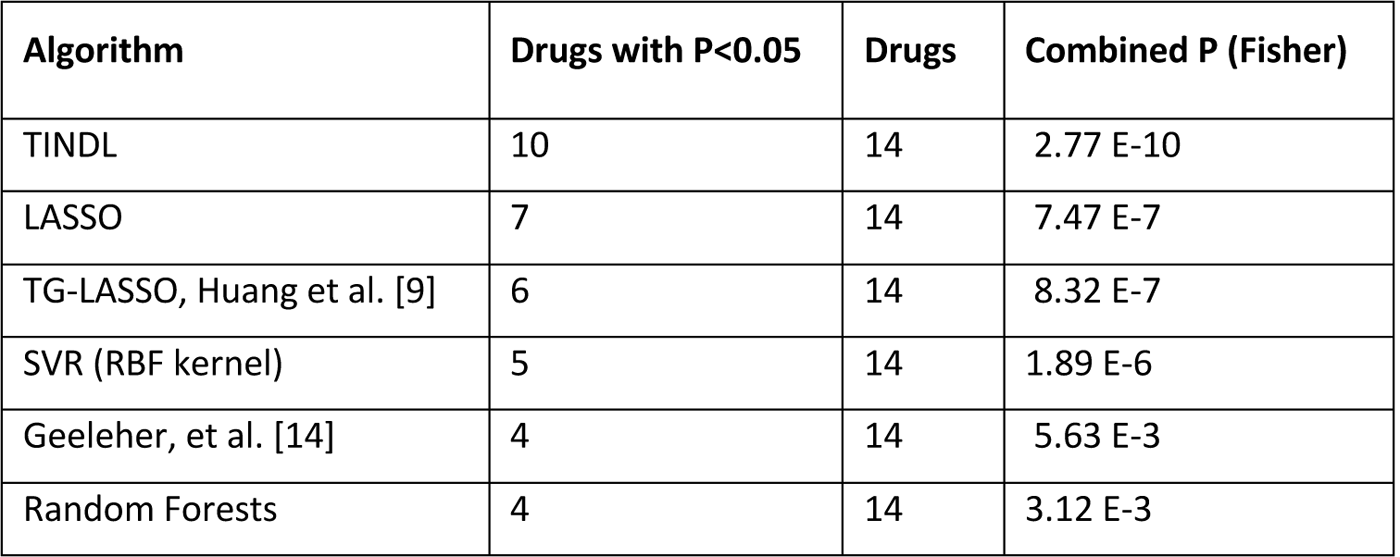
The performance of different computational models in predicting CDR of TCGA samples using models completely trained on preclinical GDSC CCLs. The first column shows the algorithm. The second column shows the number of drugs for which a one-sided Mann-Whitney U test showed a significant p-value. The third column shows the total number of drugs used for evaluation, and the fourth column shows the combined p-value (combined over all 14 drugs using Fisher’s method).

As shown in table 2, TINDL can distinguish between sensitive and resistant patients for 10 (out of 14) drugs (with a combined p-value of 2.77E-10 for all drugs), while the second-best method in this table can only distinguish between sensitive and resistant patients for 7 drugs. Similar to our previous study [9], we also observed that Lasso and its variation, TG-Lasso, perform reasonably well (when considering all drugs), while Support Vector Regression and Random Forests did not perform as well.

As discussed earlier, one of the major challenges in predicting the CDR of cancer patients based on ML models trained on preclinical CCLs is the statistical differences between these samples. To assess the performance of TINDL against other DL models that explicitly try to remove these statistical differences, we considered three alternative methods. The first method (referred to as “ComBat-DL”) utilizes ComBat [15] as a pre-processing step to remove the statistical discrepancy between CCLs and tumour samples. ComBat [15] is a popular method for removing “batch effects” in gene expression datasets and has been widely used for drug response prediction [9, 14, 19] and other applications [20, 21]. The ComBat-transformed GEx profiles are then used in a DL architecture similar to TINDL for a fair comparison. The second and third methods (called “DANN-DL” and “ADDA-DL” henceforth) are based on DANN [16] and ADDA [17], two domain adaptation techniques that were originally developed for image processing. Instead of adapting the GEx input features, these methods adjust the latent feature representations learned by the encoder. DANN uses adversarial neural networks to create a shared latent feature space between the datasets. ADDA, on the other hand, is a unidirectional domain adaptation approach that builds over a pre-trained predictor and tries to adapt the first few layers of the neural network such that the target dataset’s latent feature representation aligns with that of the source dataset. We trained models of these methods with a similar architecture to that of TINDL, with the exception of the discriminators, which are specific to ADDA and DANN and are used for domain adaptation. The details of these methods, including their architecture and training procedure are provided in Methods and Supplementary Methods. Table 3 and Supplementary Table S2 show the performance of these DL-based approaches. These results show that in all three cases, only for 7 (out of 14) drugs the predicted normalized ln(IC5) of sensitive patients is significantly smaller than resistant patients.

**Table 3:**
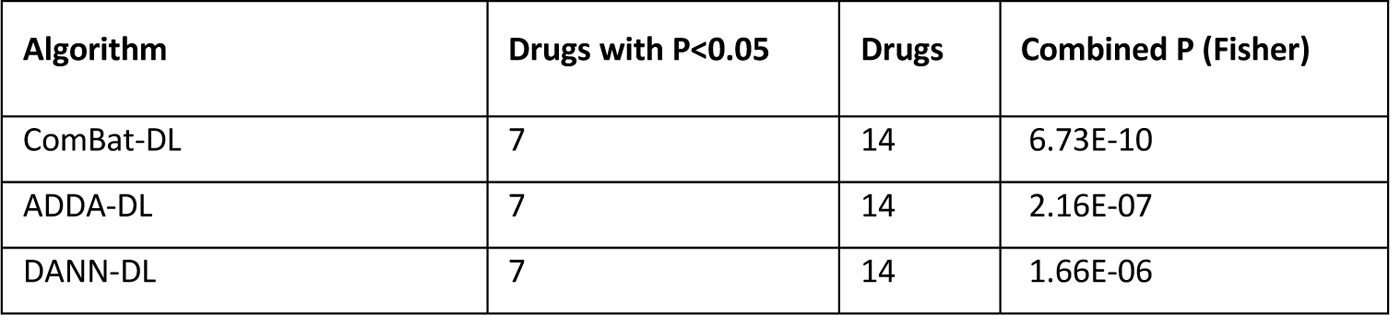
The performance of three deep learning-based methods that explicitly try to remove discrepancy between preclinical training and clinical test datasets. The first column shows the name of the algorithm. The second column shows the number of drugs for which a one-sided Mann-Whitney U test showed a significant p-value. The third column shows the total number of drugs used for evaluation, and the fourth column shows the combined p-value (combined over all 14 drugs using Fisher’s method). To ensure a fair comparison, a similar architecture to TINDL was used for all these methods. Additionally, each model was trained using ten random initializations and the mean aggregate of these predictions were used for calculating the p-values.

| Algorithm | Drugs with $P < 0.05$ | Drugs | Combined P (Fisher) |
| --- | --- | --- | --- |
| ComBat-DL | 7 | 14 | 6.73E-10 |
| ADDA-DL | 7 | 14 | 2.16E-07 |
| DANN-DL | 7 | 14 | 1.66E-06 |

To assess the superior performance of TINDL compared to these DL-based models that use an architecture similar to TINDL, we assessed their ability in removing the discrepancy between preclinical and clinical samples. For this purpose, we assessed the distance of clinical samples and preclinical samples for each method and each drug (see Methods for details of calculating distances). Since methods that use domain adaptation do not modify the input features, but rather seek to remove the domain discrepancies in the latent space (the output of the encoder), we used the learned representation of each sample in the latent space for all methods. Using a one-sided Wilcoxon signed rank test, we observed that TINDL’s learned representations for clinical samples have a significantly smaller average distance to preclinical samples compared to ComBat-DL (p = 6.10E-5), ADDA-DL (p = 4.27E-4), and DANN-DL (p = 6.10E-5), for all drugs (Figure 3A). The effectiveness of tissue-informed normalization of TINDL in removing the statistical discrepancy between the preclinical and clinical embeddings can also be visually observed using principal component analysis and UMAP plots of the representations learned by each method (Figure 3B and Supplementary Figures S1-S4).

**Figure 3:**
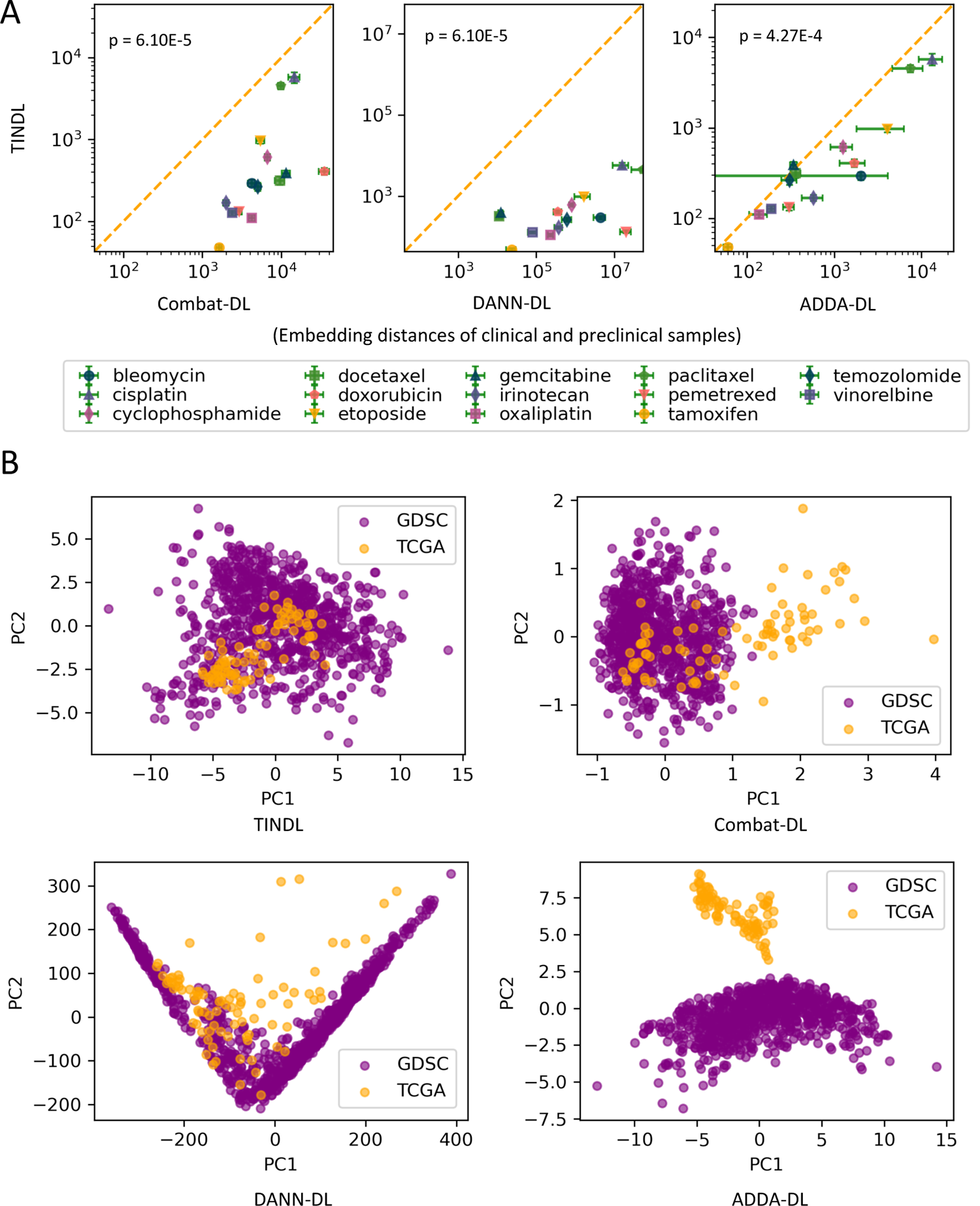
Evaluation of the embeddings used by TINDL and other deep learning methods used for prediction of drug response. A) The scatter plots compare the distance between preclinical samples and clinical samples in the embedding space for each drug. Each point in the scatter plot corresponds to a different drug. The p-values are calculated using a one-sided Wilcoxon signed rank test. The error bars show the 95% confidence intervals and are calculated based on ten runs of each method with random initializations. B) The PCA analysis of the embeddings used by each method to predict the response to etoposide. Visually, the TCGA samples are better mixed (i.e. are not easily separable) with GDSC samples in TINDL compared to other methods.

Next, we sought to determine whether the latent space representation similarity has an influence on drug response prediction performance of TINDL across different drugs. We observed a negative Spearman’s rank correlation (r = −0.17, p = 3.93 E-2) between the aforementioned distances and the AUROC of prediction for different drugs. In particular, tamoxifen that had the highest AUROC (Supplementary Table S2, AUROC=0.92), also had the smallest average distance between clinical and pre-clinical representations of its samples among all drugs in TINDL. These results further support the conclusion that reducing the discrepancy between the statistical characteristics of clinical and preclinical samples plays an important role in the success of TINDL in the prediction of CDR.

### TINDL identifies biomarkers of drug sensitivity

We used TINDL (Figure 1B) to assign a score to the contribution of each gene in the trained model (see Methods for details). Supplementary Figure S5 shows the distribution of these scores for each drug. To identify the threshold below which the contribution of the genes to the predictive model is small, we used a method called kneedle [22], which systematically determines this threshold for each drug based on the distribution of the scores. This method identified between 64 (for pemetrexed) to 243 (for bleomycin) genes, depending on the drug. The ranked list of genes identified by TINDL that pass this drug-specific threshold are provided in Supplementary Table S4.

Next, we sought to determine whether the identified genes are drug specific. To this end, we calculated the Jaccard similarity coefficient of drug pairs (Figure 4A). The results revealed a high degree of drug specificity with the average Jaccard similarity coefficient for all drugs equal to only 0.027. However, some genes were implicated for multiple drugs (Figure 4B and Supplementary Table S5). In particular, SLFN11 was implicated for nine drugs and was the top contributor for bleomycin, cisplatin, doxorubicin, etoposide, gemcitabine, and irinotecan, and the top third contributor for oxaliplatin. SLFN11 (Schlafen family member 11) is a putative DNA/RNA helicase that is recruited to the stressed replication fork and inhibits DNA replication. DNA replication is one of the fundamental biological processes in which dysregulation can cause genome instability [23]. This instability is one of the hallmarks of cancer and confers genetic diversity during tumorigenesis [24, 25]. Various studies have shown that the expression of this gene sensitizes cancer cells to many chemotherapeutic agents including cisplatin, oxaliplatin, irinotecan, gemcitabine, doxorubicin, and etoposide [26–30]. Epigenetically mediated suppression of SLFN11 via EZH2 contributes to acquired chemotherapy resistance, one that can be prevented and/or actively remodeled through targeting EZH2 [31]. Several potent and selective EZH2 inhibitors are now in different stages of clinical development with promising safety profile, including phase II (Epizyme) and phase I (Constellation, GSK) trials in multiple solid tumor and hematological indications. Our data supports the notion that the combination of downregulating SLFN11 via EZH2 inhibitor with chemotherapeutic reagents should be considered in multiple cancer types [32, 33].

**Figure 4:**
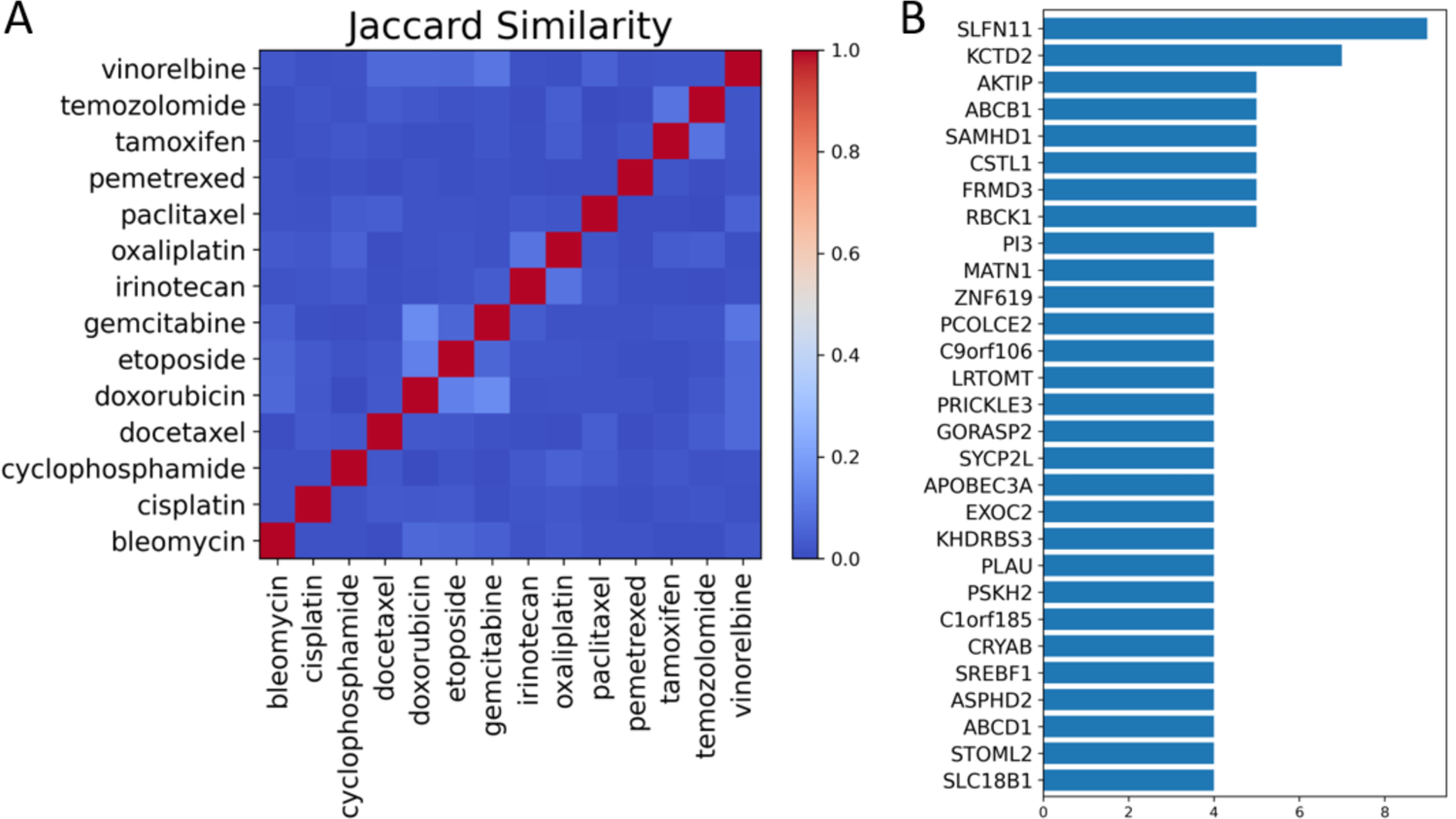
Genes identified by TINDL for different drugs. A) Heatmap of the Jaccard similarity of the selected top genes in the 14 drugs. B) Number of drugs in which the genes were identified as a top gene. Only genes that were present in the top genes of at least four drugs are included.

To better understand the functional characteristics of genes implicated by TINDL for multiple drugs, we used KnowEnG’s gene set characterization pipeline [34] to identify pathways associated with 29 genes identified by TINDL for at least 4 drugs (Figure 4B). This pipeline enables identification of associated pathways while incorporating interactions among genes and their protein products through network-guided analysis. The results (Supplementary Table S5) implicated five pathways, which included “Regulation of toll-like receptor signaling pathway”, “Alpha-synuclein signaling”, “Arf6 trafficking events”, “Insulin Pathway”, and “RalA downstream regulated genes”. Innate immune receptors such as toll-like receptors (TLRs) are responsible for recognizing molecular patterns associated with pathogens and provide critical molecular links between innate cells and adaptive immune responses. Engagement of TLRs on dendritic cells (DCs) promotes cross-talk between the innate and the adoptive immune system, maturation and migration of DCs into lymph nodes leading to activation, proliferation and survival of tumour antigen-specific naïve CD4+ and CD8+ T cells [35]. Tumour cells themselves do not express molecules which would induce DC maturation, thus application of TLR agonists is an important element of immunotherapy protocols aiming T cell activation [36]. In addition, TLR agonists have been proposed as adjuvants for cancer vaccines [37]. TLR3 agonist as an adjuvant with conventional chemotherapy can break tolerogenic or immunosuppressive effects generated by the tumour and drive T cell responses and tumor rejection [38–41].

Alpha-synuclein (α-syn) is a neuronal protein responsible for regulating synaptic vesicle trafficking. α-syn is frequently expressed in various brain tumours and melanoma [42, 43] and its upregulation has been linked to aggressive phenotypes of meningiomas [44]. Moreover, loss of α-syn results in dysregulation of iron metabolism and suppression of melanoma tumour growth [45]. Oncogenic activation of synuclein contributes to the cancer development by promoting tumor cell survival via activation of JNK/caspase apoptosis pathway and ERK and by providing resistance to certain chemotherapeutic drugs [46, 47], suggesting synuclein as a new therapeutic target for future treatment to overcome resistance to certain chemotherapeutic. ARF6 (ADP-ribosylation factor 6) governs the trafficking of bioactive cargos to tumor-derived microvesicles (TMVs) which comprise a class of extracellular vesicles released from tumor cells that facilitate communication between the tumor and the surrounding microenvironment [48]. Invasive tumor cells shed TMVs containing bioactive cargo and utilize TMVs to degrade extracellular matrix during cell invasion [49]. Indeed, several studies have suggested a correlation between expression of ARF6 and invasion and metastasis of multiple cancers [50–52], suggesting that antagonistic ARF6 signaling can dictate TMV shedding and the overall mode of invasion. Insulin, a signaling molecule that controls systemic metabolic homeostasis, can be seen as enabling tumour development by providing a mechanism for PI3K activation and enhanced glucose uptake [53–58] and plays a role in cytotoxic therapy response [59]. RalA (RAS Like Proto-Oncogene A) is a member of the Ral family, and the RalA pathway contributes to anchorage independent growth, tumorigenicity, migration and metastasis [60–64]. In conclusion, the link between genes implicated for multiple drugs and the pathways above that play different roles in cancer may point to shared mechanisms of action among different anti-cancer drugs. We also performed a similar pathway enrichment analysis for genes implicated for each drug separately and the results are provided in Supplementary Table S6.

### Functional validations confirm the role of TINDL-identified genes in response to tamoxifen

We sought to evaluate the drug response predictive ability of top identified genes by TINDL, both computationally and experimentally. We focused on tamoxifen due to the good prediction performance of TINDL for this drug (AUROC=0.92, p=1.14E-3 for Mann Whitney U test). First, using only top implicated genes for this drug (n = 136 based on the threshold identified by kneedle), we observed a consistently high value of AUROC and a significant Mann-Whitney U test p-value (Figure 5A, AUROC = 0.89, p =2.32 E-3). Next, we reduced the number of genes for the model to only top twenty and observed that AUROC remains high even with this small number of genes (Figure 5A, AUROC = 0.90, p = 1.65 E-3). This shows that even a small panel of twenty genes can be used to predict the CDR of this drug, suggesting potential clinical applications in precision medicine for these small panels of genes.

**Figure 5:**
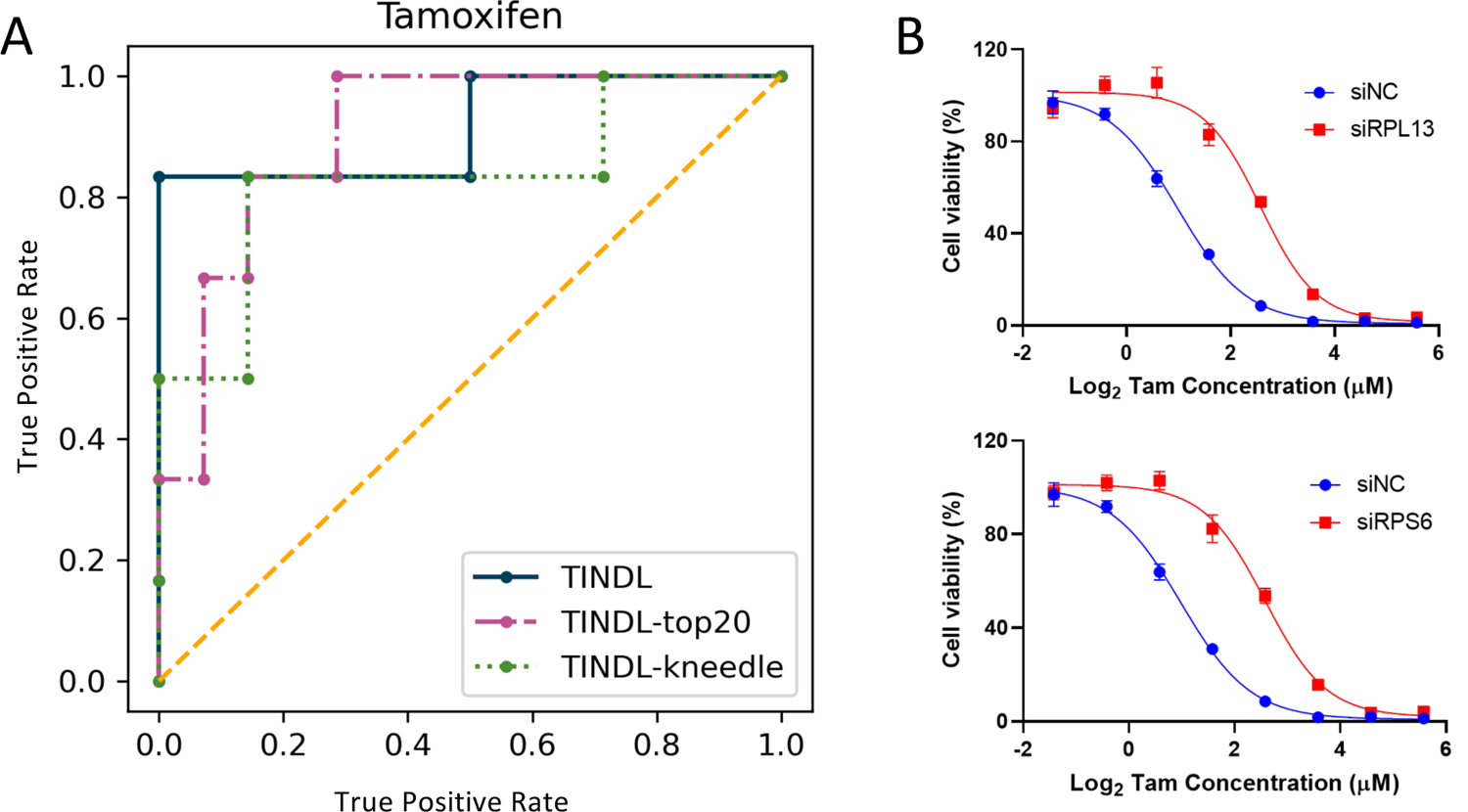
Top genes identified for tamoxifen and their functional validation. A) The ROC curve for tamoxifen, when different number of genes are used for CDR prediction. TINDL utilizes the GEx values of all genes (AUROC = 0.92), while TINDL-top20 (AUROC = 0.90) and TINDL-kneedle (AUROC = 0.83) assign a value of 0 to all genes except for top 20 and top genes identified by kneedle, respectively. B) Tamoxifen dose-response curves corresponding to the siRNA knockdown of RPS6 and RPL13 in MCF7 cells. The dose response curves for all genes are provided in Supplementary Figure S6. The p-values are calculated using an extra sum-of-squares F test.

Next, we set out to determine whether genes identified by TINDL as predictive of tamoxifen response could be associated *in-vitro* to significant changes in drug sensitivity. We selected 10 genes identified by TINDL, which included the top 9 ranked genes (RPP25, EMP1, EXTL3, EXOC2, NUP37, RPL13, WBP2NL, RPS6, GBP1) as well as the gene ranked as 19 (JAK2), due to its involvement with the type II interferon signaling pathway, an important pathway in cancer [65]. We used estrogen receptor positive breast cancer cell line, MCF7, since tamoxifen has most often been used as the treatment for estrogen receptor positive breast cancer patients in general and 85% of patients in our test dataset for this drug corresponded to breast cancer. We measured the dose-response values of tamoxifen in MCF7 cell line for these ten genes using Cyquant assay which provides an accurate measure of cell number based on DNA content [66–68]. We defined “significance” as a gene knockdown with a significant change in apparent IC50 in comparison with a negative control siRNA. Knockdown of all ten genes with specific siRNAs had a significant effect on tamoxifen sensitivity in MCF7 cell line (p<0.0001, extra sum-of-squares F test), validating 100% of tested genes in this cell line (Figure 5B, Table 4, Supplementary Figure S6). Taken together, through the functional validation in estrogen receptor positive breast cancer cells, we found that the expression of ten genes, RPP25, EMP1, EXTL3, EXOC2, NUP37, RPL13, WBP2NL, RPS6, GBP1, and JAK2, were involved in tamoxifen-induced response. The percentage of variation in the IC50 of breast cancer cells that was explained by the variation of these ten genes’ expression is provided in Table 4.

**Table 4:** The result of siRNA gene knockdown experiments in MCF7 cell line for 10 genes identified by TINDL for tamoxifen. The p-values are calculated using an extra sum-of-squares F test. Genes are sorted based on their rank by TINDL.

| Gene | Rank by TINDL | MCF7 |  |
| --- | --- | --- | --- |
|  |  | p-value | Change in IC50 |
| RPP25 | 1 | <0.0001 | 146% |
| EMP1 | 2 | <0.0001 | 69% |
| EXTL3 | 3 | <0.0001 | 101% |
| EXOC2 | 4 | <0.0001 | 89% |
| NUP37 | 5 | <0.0001 | 83% |
| RPL13 | 6 | <0.0001 | 201% |
| WBP2NL | 7 | <0.0001 | 113% |
| RPS6 | 8 | <0.0001 | 202% |
| GBP1 | 9 | <0.0001 | 113% |
| JAK2 | 19 | <0.0001 | 134% |

## DISCUSSION

Predicting the response of an individual to cancer treatments and identification of predictive biomarkers of drug sensitivity are two major goals of individualized medicine. Computational models that can achieve these goals based on preclinical *in-vitro* data can make a significant impact, due to the significant ease of preclinical data generation and data collection compared to clinical samples. This is particularly important for newly developed or newly approved drugs, for which clinical samples may be very limited or non-existent. However, the biological and statistical differences between cancer cell lines and patient tumours, make this task challenging. In a recent study [9], we assessed the ability of a wide range of machine learning models trained on preclinical CCLs, including those that incorporate auxiliary information such as gene interaction networks, in predicting the CDR of cancer patients. Our analysis confirmed the difficulty of this task and emphasized the importance of carefully designing advanced computational techniques.

In this study, we developed TINDL, and showed substantial improvement compared to state-of-the-art machine learning models (based on both traditional and deep learning techniques) (Figure 1). Our results showed the importance of removing the statistical discrepancies between preclinical and clinical samples, as well as incorporating the cancer type and tissue of origin of the tumour samples. TINDL is not simply a drug response predictor, but rather allows identification of most predictive biomarkers for each drug. The biomarkers identified by multiple drugs (Figure 4B) suggested important genes and signaling pathways that may play important roles in the mechanism of action of different drugs in cancer. Many genes identified during our study have been reported to have altered levels of expression in response to a given drug, especially SLFN11 for multiple chemotherapies [26–30], SALL4 for cisplatin [69], ABCB1 for taxane and doxorubicin [70, 71], PIGB for gemcitabine [72], and BAX to oxaliplatin [73]. These results suggest that our preclinical-to-clinical model could generate biologically relevant candidate genes and pathways for understanding mechanisms underlying drug resistance, and may offer additional combinational therapeutic strategies to overcome certain drug resistance.

Focusing on tamoxifen, we were able to show that only a small panel of 20 genes can preserve the predictive performance of TINDL for this drug (Figure 5A). Moreover, functional validation of 10 of these genes identified by TINDL using siRNA knockdown performed with MCF-7 estrogen receptor positive breast cancer cells, confirmed the direct role of these genes in response to tamoxifen (Figure 5B and Supplementary Figure S6). These results suggest that, like many complex traits, response to tamoxifen also involves multiple genes in different pathways. In addition, these results provided us with new insights into novel mechanisms in tamoxifen response. For example, among these ten genes, RPS6 is the canonical substrate of S6 kinase (S6K), which is activated by integrin engagement and inactivated by detachment. Abnormal expression of RPS6 has been indicated as a critical trigger for detachment-induced keratinization related to breast cancer development [74]. Indeed, the prognostic value of RPS6 was assessed by Kaplan-Maier Plotter analysis of gene expression data from estrogen receptor positive/HER2 negative breast tumor samples of 686 patients. High expression of RPS6 was associated with better relapse-free survival (RFS) in this cohort of patients (Supplementary Figure S7A). Decreased phosphorylation of RPS6 was previously observed in tamoxifen resistant breast cancer cells compared to parental cells [75]. However, to the best of our knowledge, no previous study has linked RPS6 to tamoxifen sensitivity. The fact that we found that RPS6 expression can predict tamoxifen sensitivity and that knockdown of RPS6 desensitized breast cancer cells to tamoxifen exposure by two folds suggests a potential role for RPS6 in the estrogen response pathway, in addition to its role as a protein synthesis regulator. In addition to its prognostic value, further analysis revealed that high mRNA expression of RPS6 was also remarkably associated with prolonged RFS in tamoxifen treated patients (Supplementary Figure S6B). This hypothesis will need to be tested further in future experiments. The second gene that influenced tamoxifen response the most was RPL13, also known as “Ribosomal Protein L13”. RPL13 is a component of the 60S ribosomal subunit that expressed at significantly higher levels in benign breast lesions than in breast carcinomas [76], however, to the best of our knowledge, no previous study has linked RPL13 to estrogen signaling or tamoxifen response. Kaplan-Meier analysis revealed that patients with high expression of RPL13 had a significantly longer RFS than those with low RPL13 expression (Supplementary Figure S7C). Our observations here suggest an important role of RPL13 expression level in predicting tamoxifen sensitivity, and could help identify additional drug targets or treatment options to overcome tamoxifen resistance.

Our analysis suggested that TINDL performs better than DL-based domain adaptation techniques in removing the discrepancies between the preclinical and clinical samples. However, these domain adaptation techniques were originally developed for the task of analyzing images. We posit that novel domain adaptation techniques may be able to overcome the shortcoming of current techniques and improve the results. However, such methods need to be carefully designed for the analysis of gene expression data and must take into account biological factors that influence the response of cancer patients to different drugs. In addition, including information on the cancer type or even subtypes of each cancer may be necessary to achieve better results.

Another important consideration is that due to the limitation of CCLs in mimicking patient tumours (e.g., their growth in 2D environment, being more homogenous than tumours, and not being able to capture the effect of tumour microenvironment, etc.), computational models trained on CCLs are limited in their ability to predict CDR of cancer patients, even if they remove the statistical discrepancies of the training and test sets. As a result, availability of large datasets, pertaining to better models of cancer, such as patient-derived organoids or xenografts play an important role in improving the predictive ability of computational models.

In this study, our focus was on models trained only on gene expression profiles of samples. However, a multi-omics approach that incorporates different molecular characteristics of samples may provide a more complete understanding of the mechanisms of drug response in cancer. Such models, however, need to be carefully designed to avoid over-fitting due to the additional number of features. Another limitation of this study was that all the computational models were trained on CCLs and their response to single drugs. However, some of the patients in the TCGA dataset have received multiple drugs in the course of their treatment, which we had to include in the analysis due to the small number of samples with known CDR. In such cases, any computational model trained on single drugs can only provide an approximation. To improve the prediction performance in such cases, a computational model must also consider the synergistic and antagonistic effects of the drugs. Recent large publicly available datasets such as DrugComb [77] and DrugCombDB [78] that contain response of different cell lines to pairs of drugs provide an opportunity for developing such methods, a direction that we will pursue in the future.

## METHODS

### Datasets

We used the publicly available data from GDSC and TCGA for training and testing, respectively. For training data, we used the RMA-normalized gene expression data in GDSC, which contains 15650 genes and 958 unique cell lines. For the test data, we used RNAseq (in FPKM) from primary tumors in TCGA, which we transformed using log(FPKM+0.1). We z-score normalized the gene expression data as well as the ln(IC50) values. We obtained clinical drug response of cancer patients from the supplementary file of Ding et al. [10]. Since the number of samples with known drug response in TCGA is relatively small, in our analysis we also included samples that have received multiple drugs in their course of treatment. We only focused on drugs which are common to both datasets and have at least 20 samples with known CDR in TCGA. We used a tissue-informed normalization, which is detailed below. Furthermore, we re-categorized the clinical drug responses to sensitive (corresponding to complete and partial response) and resistant (corresponding to stable disease and clinically progressive disease). Details on sample counts and tissue types per drug are in Supplementary Table S1.

### Tissue-informed Normalization

TINDL trains a separate model for each drug. Each model performs a separate normalization on the GEx profiles of test samples from TCGA to account for the cancer type and tissue of origin of the samples. First, for each drug *D* the set of tissues/cancer types to which this drug is administered in the TCGA samples is identified (referred to as *T*_D_). All samples corresponding to *T*_D_ (excluding those used in the test set) are collected from TCGA, forming the unlabeled dataset. Then, the gene-wise mean (μ_TD_) and standard deviation (σ_TD_) of these unlabeled samples are calculated and used to normalize labeled test samples corresponding to drug *D*. More specifically, for a gene *i* of an arbitrary sample in the test set, the normalized value *x*_i_ would be:

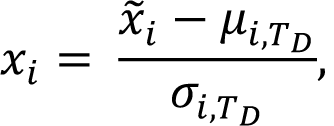

where 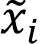 is the log-transformed expression for gene *i* of that sample. The test samples are then used as input to the trained model to predict the normalized ln(IC50)s, which were compared to the actual CDR categories for evaluation.

### TINDL Architecture, hyperparameter selection and training

We used grid-search and 5-fold cross validation to select the number of epochs, batch size and learning rate of all our DL-based models (including TINDL). Specific hyperparameters chosen using this procedure for TINDL are provided in the Supplementary table S7. We only used the training data corresponding to CCLs (from GDSC) to perform the hyperparameter search. In addition to the input layer (which contained one node for each gene), we used three hidden layers with dense connections, each with 512, 256, and 128 hidden nodes, in the order of their distance to the input layer. We used a rectified linear units (ReLU) activation function and added a dropout layer with 0.2 probability of dropping out prior to the output layer.

Models were trained using mean squared error (MSE) as the loss function, and the normalized ln(IC50) as the labels. During hyperparameter tuning, models were allowed to train up to a maximum of 1000 epochs, but early stopping was applied when the model’s loss did not decrease after 30 epochs. After hyperparameter tuning, we retrained a final model using all the labeled CCL samples. We used 10 different random initializations (i.e., seeds) to ensure robustness of the results. A similar technique was used for ADDA-DL, DANN-DL, and ComBat-DL.

### Calculating contribution scores of genes

In the second phase of TINDL (Figure 1B) we used CXPlain [18] as the explainer to assign a contribution score to each gene in each sample. CXPlain is a method that attempts to provide causal explanations of a trained model’s predictions. This is achieved by training a separate model (called “explainer”) using the outputs of the trained model (called “predictor”). This method utilizes Granger’s causality [79] to evaluate the contribution of a single feature (gene in our case) by zeroing out features one by one and calculating the normalized difference of the predictor’s original error and its error when the feature is zeroed out. In our case, we define error as 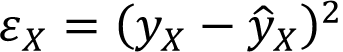, where y_X_ is the true value and ŷ_X_ is the output of the predictor for sample X = {*x*_1_, …, *x*_p_} being the number of features. Prior to training the explainer, the real contribution vectors, Ω_X_ = {ω_1_(X), …, ω_p_(X)}, are calculated for each training sample as follows:

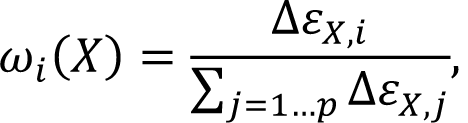

where *Δε_x,i_ = ε_X\{i}_ − ε_X_*. Here, *ε_X\{i}_* denotes the predictor’s error when given X but with feature *i* zeroed out. The explainer has an architecture such that the dimensions of the input vector X and the output vector 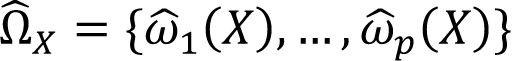 are the same. Each of the outputs correspond to the predicted contribution for the corresponding feature. The explainer is trained by minimizing the KL divergence 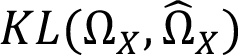 of the real contributions Ω_X_ and predicted contributions 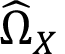 of the training set.

We used a neural network with two layers and 512 hidden units for the explainer, and used the ensemble mode, which trains 10 independent explainers and reports their median as the final contribution values. We modified the CXPlain library’s code to fit our application, which we also included in our published code. Once trained, we predicted the contribution values of each genes in each of the samples in the testing set. To obtain drug-specific gene contribution scores, we calculated the mean contribution score of each gene across all the labeled test samples for that drug and normalized it such that the largest contribution score of a drug equals 1.

### Identifying genes with highest contribution scores

After obtaining contribution scores to each gene for a drug, we sought to identify the top genes that substantially affect our model’s predictions. We sorted the genes according to their final test contribution scores and plotted a curve (Supplementary Figure S5), where the x-axis is the rank of the gene *i* and the y-axis is gene *i*’s drug-specific contribution score 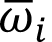. We used the kneedle algorithm [22] to identify the point of maximum curvature, called “knee”, which we then treated as the cutoff for the top genes. Kneedle relies on the idea that if one forms a line *l* from 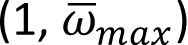 to 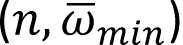 and rotate the curve around the point 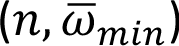, the “knee” can be approximated by the set of points in the local maxima. Among these points, the point that is farthest from the line *l* is then identified as the knee.

### Knowledge-guided Pathway Enrichment Analysis

We identified pathways associated with the top identified genes using KnowEng’s gene set characterization (GSC) pipeline [34]. We used the network-guided mode, which incorporates knowledge in the form of gene-gene interactions to augment the analysis. For the knowledge network, we selected the STRING Experimental PPI [80], which contains experimentally verified protein-protein interactions. We then proceeded with the default 50% network smoothing parameter and used the “Enrichr” pathway collection. This pipeline does not provide a p-value, but rather uses a score called “Difference Score” to implicate top pathways. Any pathway above the 0.5 threshold is considered associated with the input query set. A value above this threshold shows that the pathway has a high relevance score to the input query set (using a random walk with restarts algorithm), compared to the background [34].

### Precision at kth percentile

For each drug, we used TINDL’s predictions of ln(IC50) of the tumour samples, and identified the kth percentiles of the distribution (k ≤ 50), which we denote as t_k_. We stratified the predictions such that all predictions below t_k_ is predicted as positives (i.e. sensitive). We then calculated the precision at kth percentile as 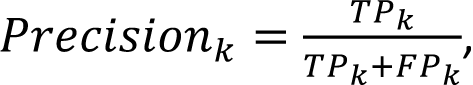 where *TP_k_* and *FP_k_* are the true positives and false positives at kth percentile, respectively.

### Baseline models

SVR, Random Forest, and Lasso Regression were all implemented using the Scikit-learn. Geeleher’s method [14] was reimplemented using Scikit-Learn and PyComBat, a python implementation of ComBat [15]. We used the available implementation of TG-Lasso [9]. All hyperparameters were tuned as described in the previous subsections except for TG-Lasso, which has its built-in hyperparameter tuning.

To ensure a fair comparison, all DL-based baseline models used a similar architecture to TINDL. Additionally, the hyperparameter tuning and training procedure was also similar to the one described above for TINDL. Below, we describe model-specific considerations. For ComBat-DL we used ComBat [15] for removing the discrepancy between TCGA and GDSC datasets. Similar to TINDL, we used both labeled and unlabeled samples of TCGA for this purpose.

ADDA-DL utilizes adversarial discriminative domain adaptation (ADDA) [17], to remove the discrepancy between TCGA and GDSC datasets. ADDA is a unidirectional domain adaptation technique, which takes a pre-trained neural network and attempts to adapt the network to the target dataset by forcing the latent feature space of the target dataset (TCGA) to be similar to that of the source dataset’s (GDSC). We used the TINDL model as the pre-trained network which we adapt through ADDA’s adversarial losses. We used the unlabeled tumour samples from the drugs target tissues during training. Details are provided in the Supplementary Methods.

DANN-DL utilizes domain adaptive neural network (DANN) [16] to remove the discrepancies between TCGA and GDSC datasets. DANN utilizes the shared latent feature space to allow the model to be used on the target dataset despite only being trained using the source dataset’s labels. This is done by incorporating a gradient-reversed discriminative loss function such that a discriminator cannot tell whether the given embedding came from the source (GDSC) or target (TCGA) datasets. Similar to ADDA-DL, we used the unlabeled tumours from the drug’s target tissues for training of the discriminator.

### Measuring distance of clinical and preclinical samples in the latent space of DL-based models

To assess the ability of each DL-based model in removing discrepancy between preclinical and clinical samples, we used pairwise Euclidean distance of samples based on their representation learned by the encoder of the DL models. Since these representations are used by the decoder to make predictions, comparing these latent representations is more meaningful than comparing input feature representations. We used Ward’s method [81] to assess the distance of preclinical samples and clinical samples, which is one of the most popular methods in assessing the distance of two groups of samples. This method, that is widely used in hierarchical clustering, has the advantage that not only analyzes the Euclidean distances of the data points, but also incorporates their variance in determining the distance of two groups of samples.

### Chemicals and reagents

Dulbecco’s minimum essential medium (DMEM) medium was purchased from Life Technologies, Inc. (Carlsbad, CA, USA). Fetal bovine serum (FBS) and charcoal-stripped FBS were from Invitrogen (Carlsbad, CA, USA). Ontarget-plus SMARTpool small interfering RNAs (siRNA) targeting RPP25, EMP1, EXTL3, EXOC2, NUP37, RPL13, WBP2NL, RPS6, GBP1, and JAK2 as well as negative control siRNA were purchased from Dharmacon (Thermo Scientific Dharmacon, Inc.). Reagents and primers for real time PCR were purchased from Qiagen (Valencia, CA, USA). 17β-estradiol (E2) and 4-hydroxytamoxifen (OH-TAM) were purchased from Sigma Aldrich (Saint Louis, MO USA).

### Cell lines

MCF-7 cell line wase obtained from American Type Culture Collection (ATCC) (Manassus, VA) in 2014 and the identities of all cell lines were confirmed by the medical genome facility at Mayo Clinic Center (Rochester MN) using short tandem repeat profiling upon receipt. MCF-7 cells were cultured in DMEM containing 10% fetal bovine serum (FBS).

### Transfection and gene silencing

Specific short interfering RNAs (siRNAs) that targeted RPP25, EMP1, EXTL3, EXOC2, NUP37, RPL13, WBP2NL, RPS6, GBP1, JAK2, and negative siRNA controls (Dharmacon, Lafayette, CO) were transfected into MCF-7 cells in 96-well plates using Lipofectamine RNAiMAX Reagent (Thermo Fisher Scientific, Waltham, MA) according to the vendor’s protocol [67, 68]. Total RNA was extracted 48 hours after transfection for RNA quantification. Specific siGENOME siRNA SMARTpool reagents against a given gene as well as a negative control, siGENOME Non-Targeting siRNA, were purchased from Dharmacon Inc. (Lafayette, CO, USA). For the purpose of drug tamoxifen response assay, cells were plated in base medium supplemented with 5% charcoal stripped FBS for 24 hours, and then cultured in FBS-free DMEM media for another 24 hours before transfection. Different treatments were started 24 hours after transfection.

### Tamoxifen sensitivity assay

Drugs were dissolved in DMSO, and aliquots of stock solutions were frozen at −80°C. Cytotoxicity assays were performed in triplicate at each drug concentration. Specifically, 4000 breast cancer cells were seeded in 96-well plates and were cultured in base media containing 5% (vol/vol) charcoal-stripped FBS for 24 hours and were subsequently cultured in FBS-free base media for another 24 hours. Cells were then transfected with either control siRNA or siRNA targeting specific gene. Twenty-four hours after transfection the media was replaced with fresh FBS-free base media and the cells were treated with 10 μL of tamoxifen at final concentrations of 0, 0.1875, 0.375, 0.75, 1.5, 3, 6, 12, 24, and 48 μM [82]. After incubation for an additional 72 hours, cytotoxicity was determined by quantification of DNA content using CYQUANT assay (#C35012, Invitrogen) following the manufacturer’s instructions [66, 83, 84]. 100μL of CyQUANT assay solution was added, and plates were incubated at 37°C for one hour, and then read in a Safire2 plate reader with filters appropriate for 480 nm excitation and 520 nm emission.

## DECLARATIONS

**Ethics approval and consent to participate:**

Not applicable.

## Consent for publication

Not applicable.

## Availability of data and materials

An implementation of TINDL in python, with appropriate documentation, is available at: https://github.com/ddhostallero/tindl. Data generated in this study are provided as supplementary files.

## Authors’ contributions

AE and JC conceived the study and designed the project. AE led the computational aspects of the study. DEH designed the algorithms, implemented the pipeline and performed the statistical analyses of the results. JC led the experimental validation of the results. LW performed the gene knockdown experiments. All authors contributed to the drafting of the manuscript and critical discussion of the results. All authors read and approved the final manuscript.

## Additional Files

**Additional File 1:**

This file contains Supplementary Methods, all supplementary figures and their captions, as well as captions of all supplementary tables. Some tables are provided as separate files.

