## Additional File 1 for "A Deep Learning Framework for Prediction of Clinical Drug Response of Cancer Patients and Identification of Drug Sensitivity Biomarkers using Preclinical Samples"

### Supplementary Methods

#### ADDA

Adversarial Discriminative Domain Adaptation (ADDA) [1] is a unidirectional domain adaptation which takes a pre-trained neural network and attempts to adapt the network to the target dataset ($T$) by forcing the latent feature space of the target datasets to be similar to that of the source dataset's ($S$). First, a model is trained on the source dataset (CCLs in our case) until it converges or a certain performance metric is achieved. We then clone this model and train only the first $n$ layers of the cloned model (here first two layers), which is called the "target encoder" ($M_t$). Samples in the target dataset would pass through $M_t$, while samples from the source dataset would use the original model's parameters, which we will call the "source encoder" ($M_s$). We then create third model, called the "discriminator" ($D$), which classifies whether the extracted features came from the source or the target dataset. This is trained using the following loss function:

$$\mathcal{L}_D = -\,\mathbb{E}_{X_s \sim S}[\log\, D_1\big(M_s(X_s)\big)] - \,\mathbb{E}_{X_t \sim T}[\log D_0(M_t(X_t))].$$

where $D_0$ and $D_1$ are the outputs of the discriminator in first and second index, respectively. This pertains to the input's predicted probability of coming from the target and source datasets and is constrained such that $D_0 = 1 - D_1$.

Oppositely, the objective of the $M_t$ is to create representations that could confuse the discriminator, thus creating an adversarial optimization with the following loss:

$$\mathcal{L}_{M_t} = -\mathbb{E}_{X_t \sim T}[\log D_1(M_t(X_t))].$$

The target encoder and the discriminator are trained alternately until convergence or the maximum number of steps are achieved.

We used the TINDL models as the pre-trained models and the first two layers as the feature extractors. The discriminator has one hidden layer with 128 neurons and two output neurons. We then selected the other hyperparameters (learning rate and number of training epochs) using 5-fold CV where we used the maximum discriminator confusion as the criterion for selecting the hyperparameter.

**DANN**

Domain Adaptive Neural Network (DANN) [2] uses the shared latent feature space to allow the model to be used on the target dataset ($T$) despite only being trained using the source dataset's ($S$) labels. The model has three components: encoder ($M$), predictor ($F$), and discriminator($D$). The predictor is the model for the main task and it is trained using the appropriate loss function (e.g. mean squared error for regression tasks, negative log-likelihood or cross entropy for classification tasks). The input of the predictor comes from the encoder, which extracts domain-invariant features from the data. The discriminator attempts to identify whether the extracted features came from the source or the target dataset. Ideally, the extracted features must have enough information for the main task, and at the same time confuse the discriminator. The model is trained using the following objective function:

$$\mathcal{L} = \mathbb{E}_{(X_s,y)\sim S}\left[\left(F\left(M(X_s)\right) - y\right)^2\right] + R_\lambda(\mathcal{L}_D).$$

Here, $\mathcal{L}_D(\cdot)$ is the classification loss of the discriminator, which given by:

$$\mathcal{L}_D = -\,\mathbb{E}_{X_s\sim S}[\log\, D_1\left(M(X_s)\right)] - \mathbb{E}_{X_t\sim T}[\log\left(D_0\left(M(X_t)\right)\right],$$

where $D_0$ and $D_1$ are the outputs of the discriminator in first and second index, respectively. This pertains to the input's predicted probability of coming from the target and source datasets and is constrained such that $D_0 = 1 - D_1$. A gradient reversal technique was used on the discriminator loss to allow end-to-end training and is denoted by $R_\lambda$ in objective function. $R_\lambda$ does not affect the network during forward propagation, but $R_\lambda$ negates the gradients during backward pass.

We used the TINDL models as the pre-trained models and the first two layers as the feature extractors. The discriminator has one hidden layer with 128 neurons and two output neurons. We then selected the other hyperparameters using 5-fold CV as described in the Methods section.

**Kaplan Meier survival analysis**

An online database Kaplan-Meier plotter (http://kmplot.com/analysis/) [3] was used to investigate the association between RPS6 and RPL13 mRNA levels and survival of estrogen receptor positive/HER2 negative breast cancer patients. The patients in datasets were classified into two groups according to auto select best cutoff (high vs. low expression). The hazard ratio (HR) with 95 % confidence intervals (CI) and log rank P value were calculated and shown on the web pages. P value of < 0.05 was considered to be statistically significant.

### Supplementary Tables

**Supplementary Table S1.** Information regarding the samples used in both training and testing sets. The table is provided as a separate file called TableS1_Data.xlsx. The first sheet corresponds to TCGA samples. Sensitive samples correspond to those with CDR of complete or partial response, while resistant samples correspond to those with progressive or stable disease. Unlabeled samples correspond to those used for tissue-informed normalization. The second sheet corresponds to GDSC samples used for training.

**Supplementary Table S2.** Performance of different models in predicting CDR. The tables are provided in a separate file called TableS2_Performance.xlsx. The first sheet compares TINDL against shallow learning (i.e. traditional machine learning) methods, while the second sheet compares its performance against deep learning methods. Top rows show p-values calculated using a one-sided Mann-Whitney U test, while the bottom rows show the AUROCs.

**Supplementary Table S3.** Precision at kth percentile of TINDL. This measure captures the ability of the model to identify sensitive patients based on different thresholds, determined based on the value of k. Detailed procedure for this calculation is provided in the Methods section.

| Drug | k = 10 | k = 20 | k = 30 | k = 40 | k = 50 |
|---|---:|---:|---:|---:|---:|
| bleomycin | 1.000 | 0.909 | 0.938 | 0.952 | 0.962 |
| cisplatin | 0.935 | 0.885 | 0.868 | 0.843 | 0.862 |
| cyclophosphamide | 1.000 | 1.000 | 1.000 | 1.000 | 0.980 |
| docetaxel | 0.455 | 0.619 | 0.645 | 0.659 | 0.647 |
| doxorubicin | 0.800 | 0.800 | 0.833 | 0.800 | 0.780 |
| etoposide | 1.000 | 0.941 | 0.960 | 0.971 | 0.976 |
| gemcitabine | 0.625 | 0.563 | 0.553 | 0.571 | 0.544 |
| irinotecan | 0.000 | 0.200 | 0.286 | 0.222 | 0.333 |
| oxaliplatin | 0.667 | 0.727 | 0.688 | 0.727 | 0.741 |
| paclitaxel | 0.688 | 0.719 | 0.771 | 0.730 | 0.759 |
| pemetrexed | 0.250 | 0.625 | 0.417 | 0.467 | 0.526 |
| tamoxifen | 1.000 | 1.000 | 1.000 | 0.875 | 0.900 |
| temozolomide | 0.200 | 0.211 | 0.172 | 0.184 | 0.167 |
| vinorelbine | 1.000 | 1.000 | 0.889 | 0.917 | 0.933 |

**Supplementary Table S4.** List of top genes identified by our pipeline. The tables are provided in a separate file called TableS4_top_genes.xlsx. Each sheet includes the ranked list of genes for a different drug. The first column in each tab is the Ensemble ID of the top genes, while the second column indicates the HGNC gene name. The third column is the contribution score of the gene.

**Supplementary Table S5.** Genes and pathways associated with multiple drugs. The tables are provided in a separate file called TableS5_MultiDrugGenes.xlsx. The first sheet contains a gene by drug table in which a value of 1 shows that the gene was implicated for the drug, while a value of 0 shows that it was not. The last column shows the total number of drugs for which each gene was implicated. The second sheet contains the results of KnowEnG's network-guided gene set characterization for 29 genes identified for at least 4 drugs. The pipeline was run with default parameters, experimentally verified protein-protein interactions from the STRING database was used for the network, and Enrichr pathways were used as target pathways. This pipeline does not provide a p-value of association, but rather uses a measure called "Difference Score" (fourth column in the second sheet) to determine associated pathways. Pathways with scores above 0.5 are considered to be associated with the input query gene set (based on a random walk with restart algorithm), when compared against the background (which captures the effect of the network).

**Supplementary Table S6.** Pathways associated with genes implicated for each drug by TINDL. The tables are provided in a separate file called TableS6_PathwayAnalysis_EachDrug.xlsx. Each sheet contains the results of KnowEnG's network-guided gene set characterization pipeline for genes identified by TINDL for each drugs. The pipeline was run with default parameters, experimentally verified protein-protein interactions from the STRING database was used for the network, and Enrichr pathways were used as target pathways. This pipeline does not provide a p-value of association, but rather uses a measure called "Difference Score" (fourth column in the second sheet) to determine associated pathways. Pathways with scores above 0.5 are considered to be associated with the input query gene set (based on a random walk with restart algorithm), when compared against the background.

**Supplementary Table S7.** Learning rates, and batch sizes, and number of epochs used in the final models. These hyperparameters were selected using 5-fold cross validations on the CCL samples.

| drug | learning rate | batch size | num. epochs |
|---|---|---|---|
| bleomycin | 0.00001 | 128 | 38 |
| cisplatin | 0.0005 | 128 | 24 |
| cyclophosphamide | 0.0001 | 128 | 6 |
| docetaxel | 0.00005 | 64 | 10 |
| doxorubicin | 0.0001 | 64 | 23 |
| etoposide | 0.0001 | 64 | 33 |
| gemcitabine | 0.00005 | 64 | 28 |
| irinotecan | 0.0001 | 64 | 21 |
| oxaliplatin | 0.00001 | 64 | 31 |
| paclitaxel | 0.0005 | 64 | 38 |
| pemetrexed | 0.00001 | 128 | 50 |
| tamoxifen | 0.00001 | 64 | 21 |
| temozolomide | 0.00001 | 128 | 39 |
| vinorelbine | 0.00005 | 128 | 8 |

**Supplementary Figures:**

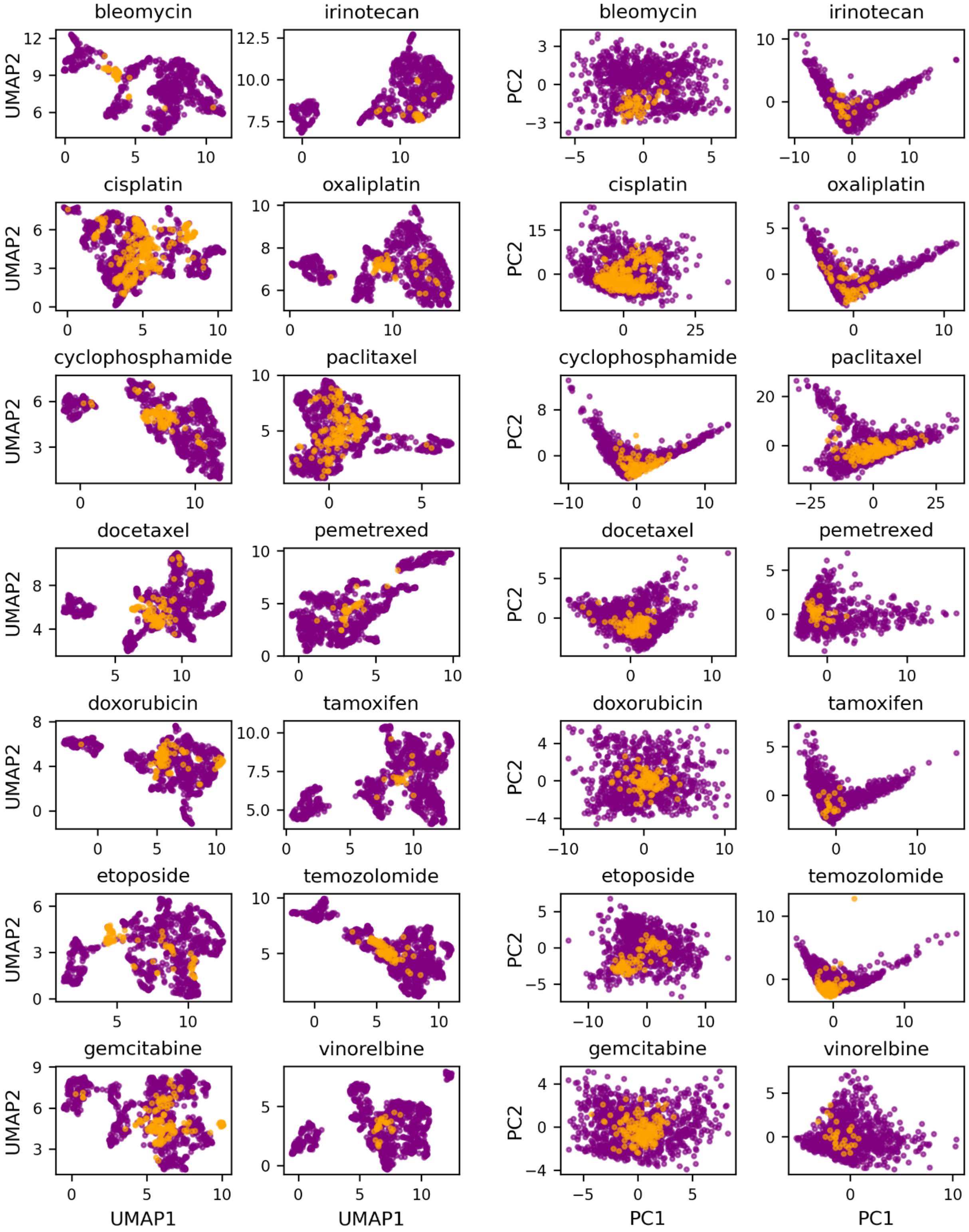


**Supplementary Figure S1**. UMAP and PCA plots of the learned latent features of different drugs by TINDL. Purple points indicate CCL samples (GDSC) and orange points indicate tumour samples (TCGA).

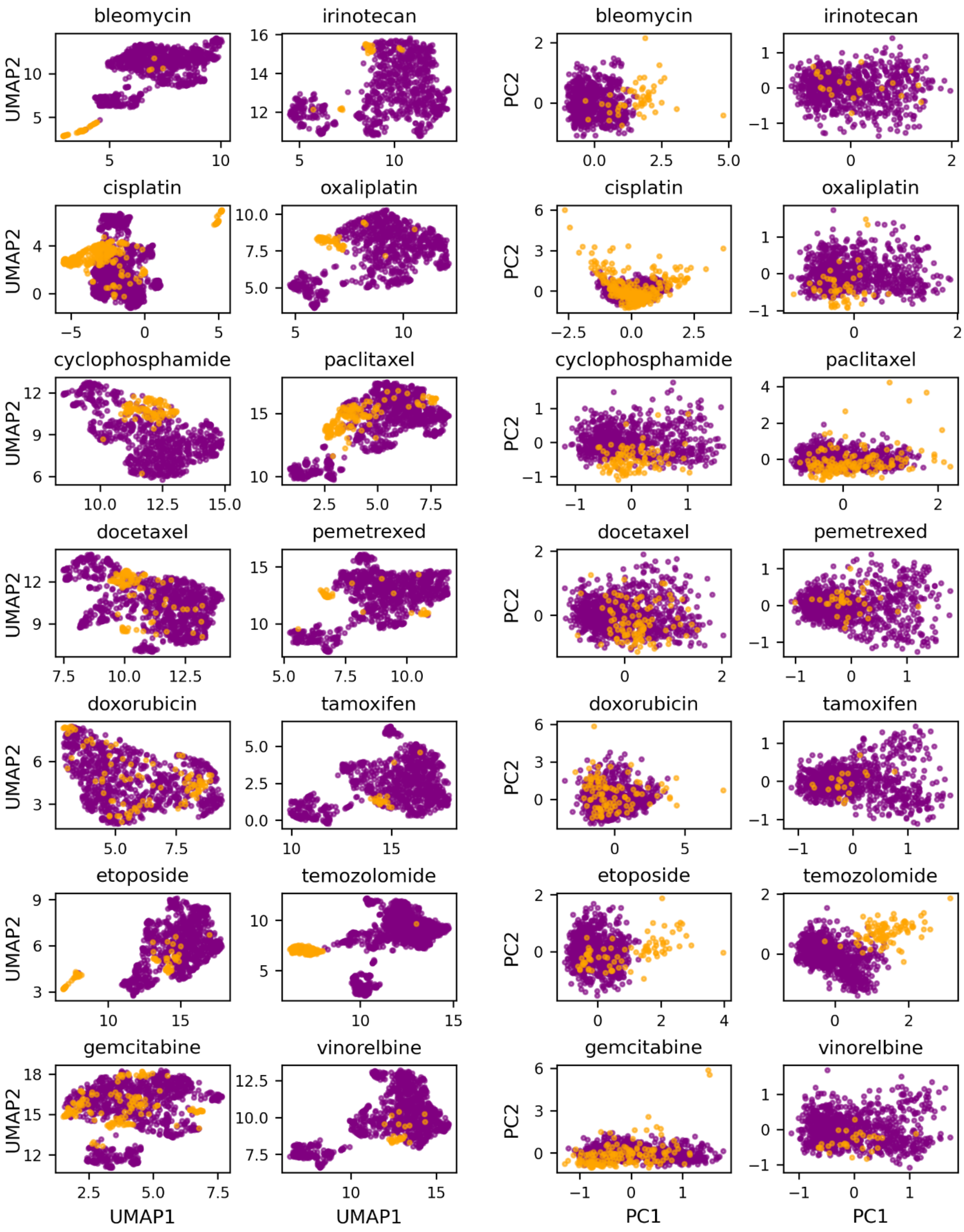


**Supplementary Figure S2**. UMAP and PCA plots of the learned latent features of different drugs by ComBat-DL. Purple points indicate CCL samples (GDSC) and orange points indicate tumour samples (TCGA).

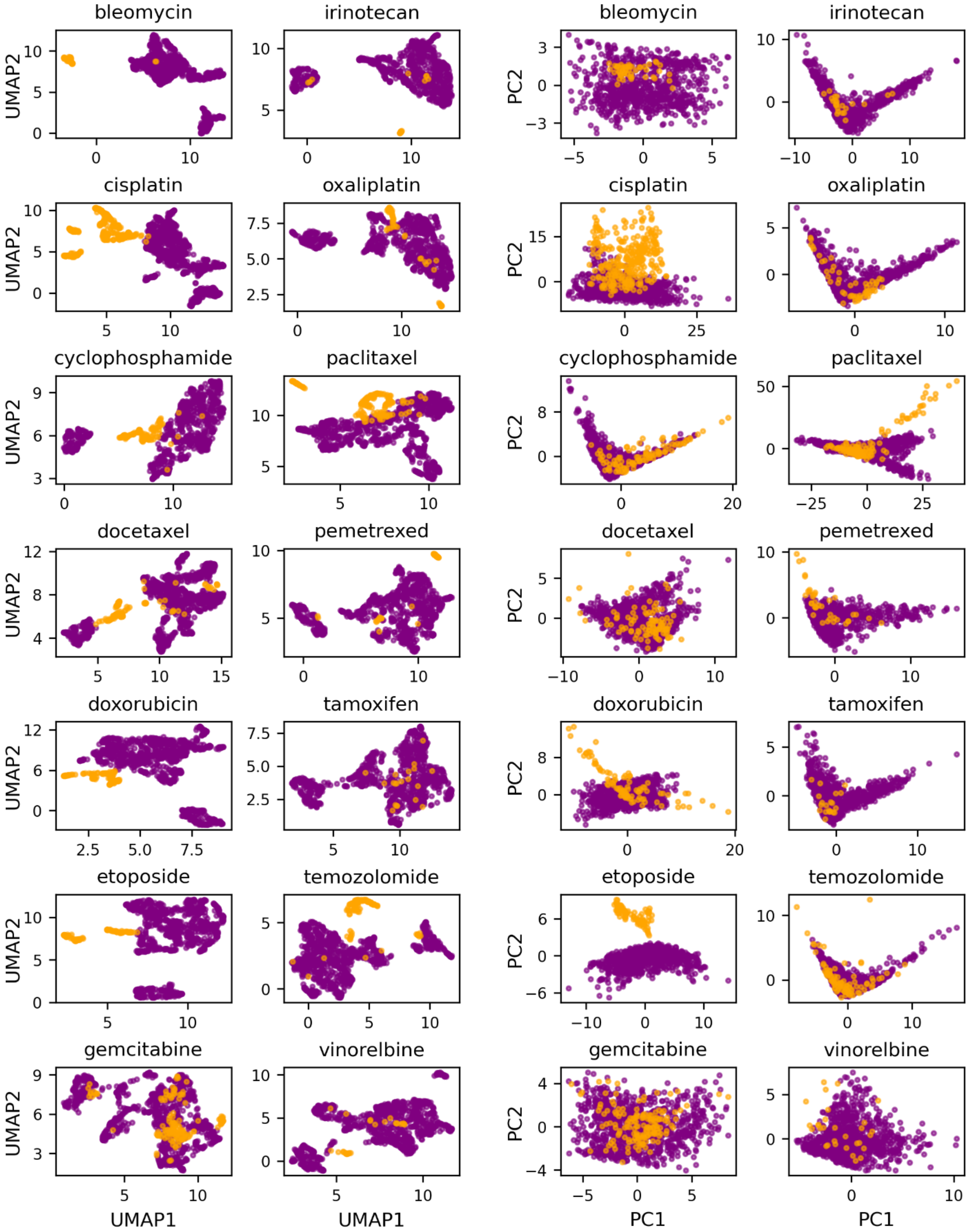


**Supplementary Figure S3**. UMAP and PCA plots of the learned latent features of different drugs by ADD-DL. Purple points indicate CCL samples (GDSC) and orange points indicate tumour samples (TCGA).

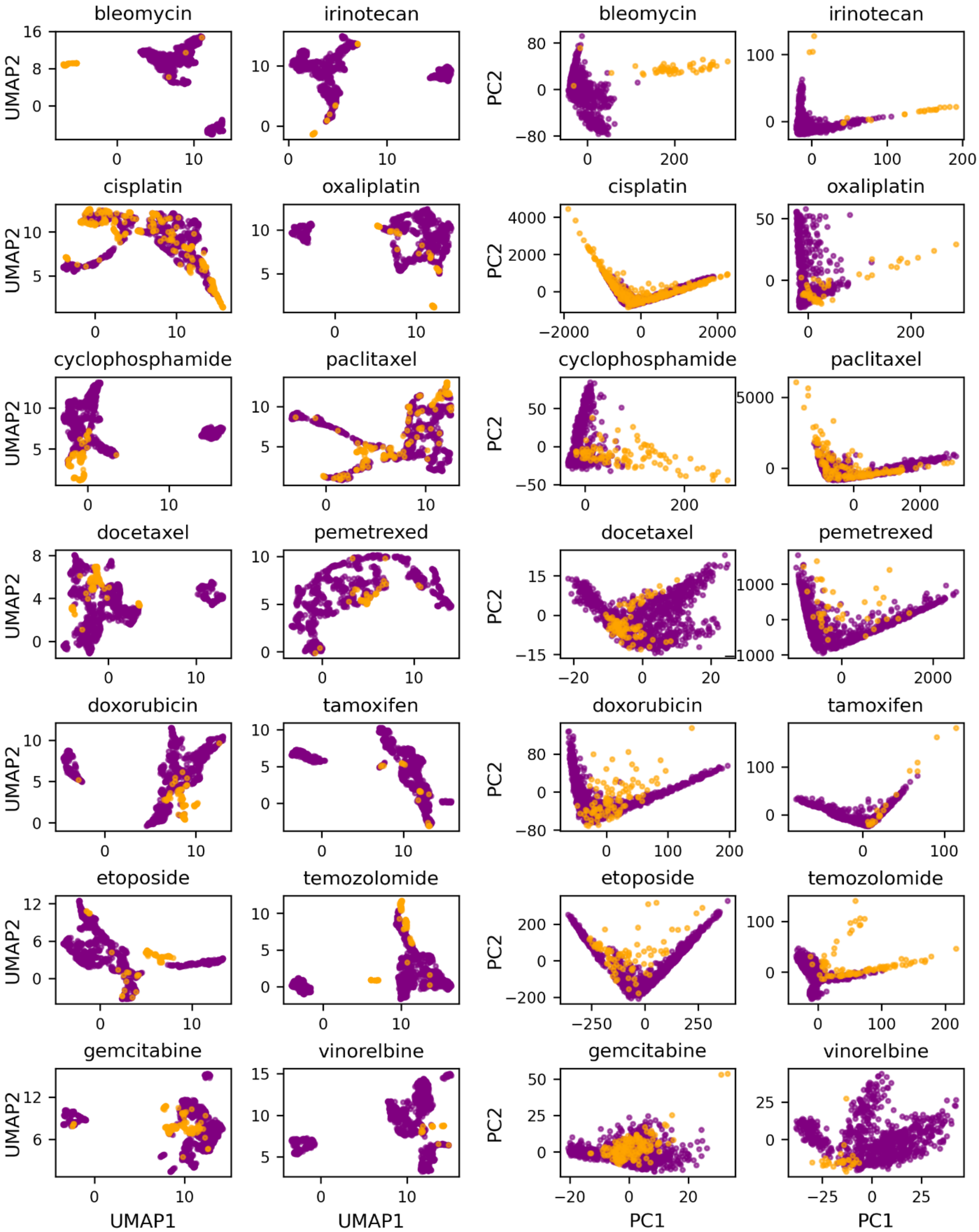


**Supplementary Figure S4**. UMAP and PCA plots of the learned latent features of different drugs by DANN-DL. Purple points indicate CCL samples (GDSC) and orange points indicate tumour samples (TCGA).

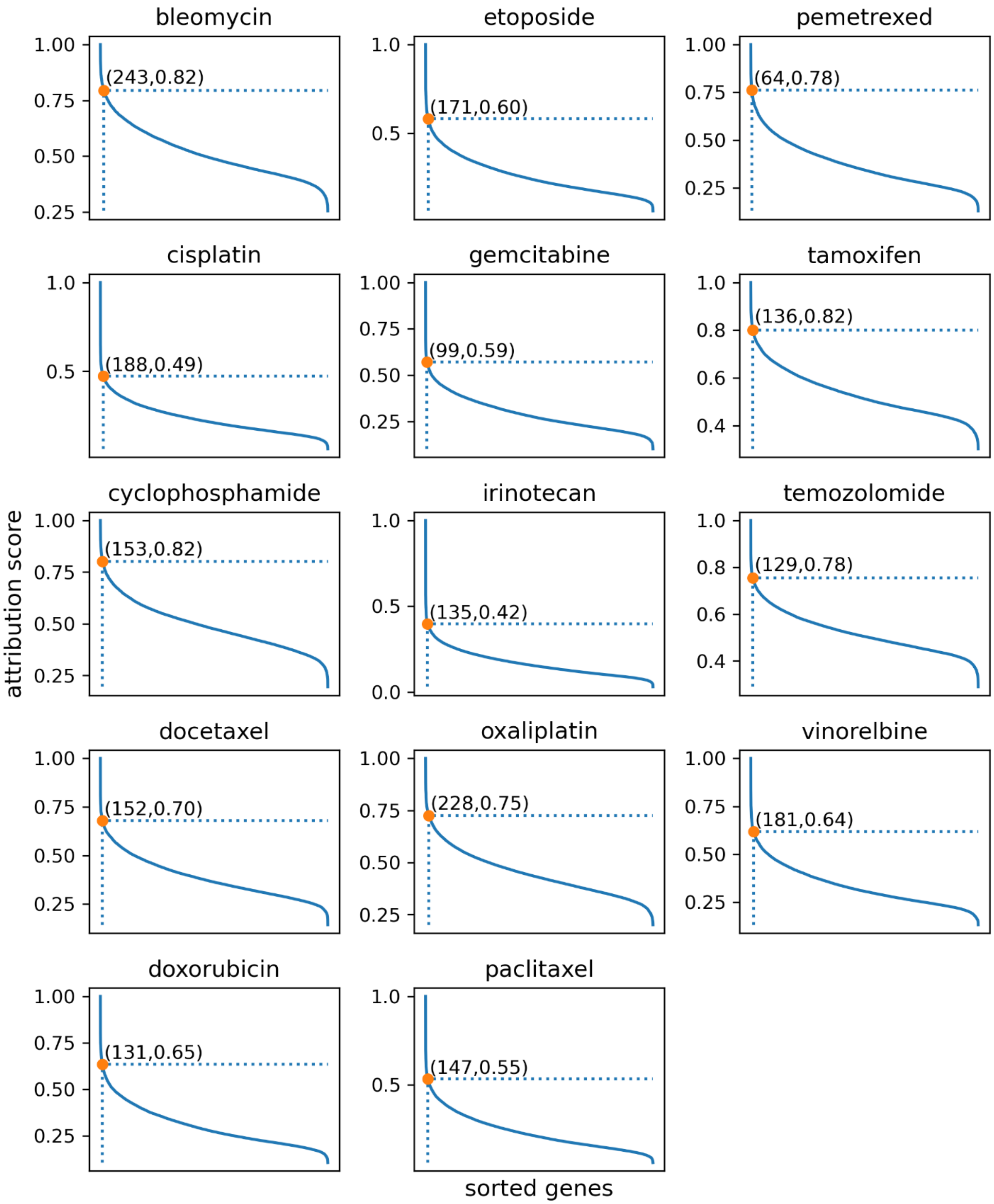


**Supplementary Figure S5**. Contribution scores of genes in the trained model of TINDL. The y-axis shows the contribution score, and the x-axis shows genes in a descending order of their score. Orange points indicate the "knees" found by kneedle algorithm [4], representing the threshold below which the contribution of genes to the trained model is small. The numbers in the brackets show (number of genes above threshold, threshold).

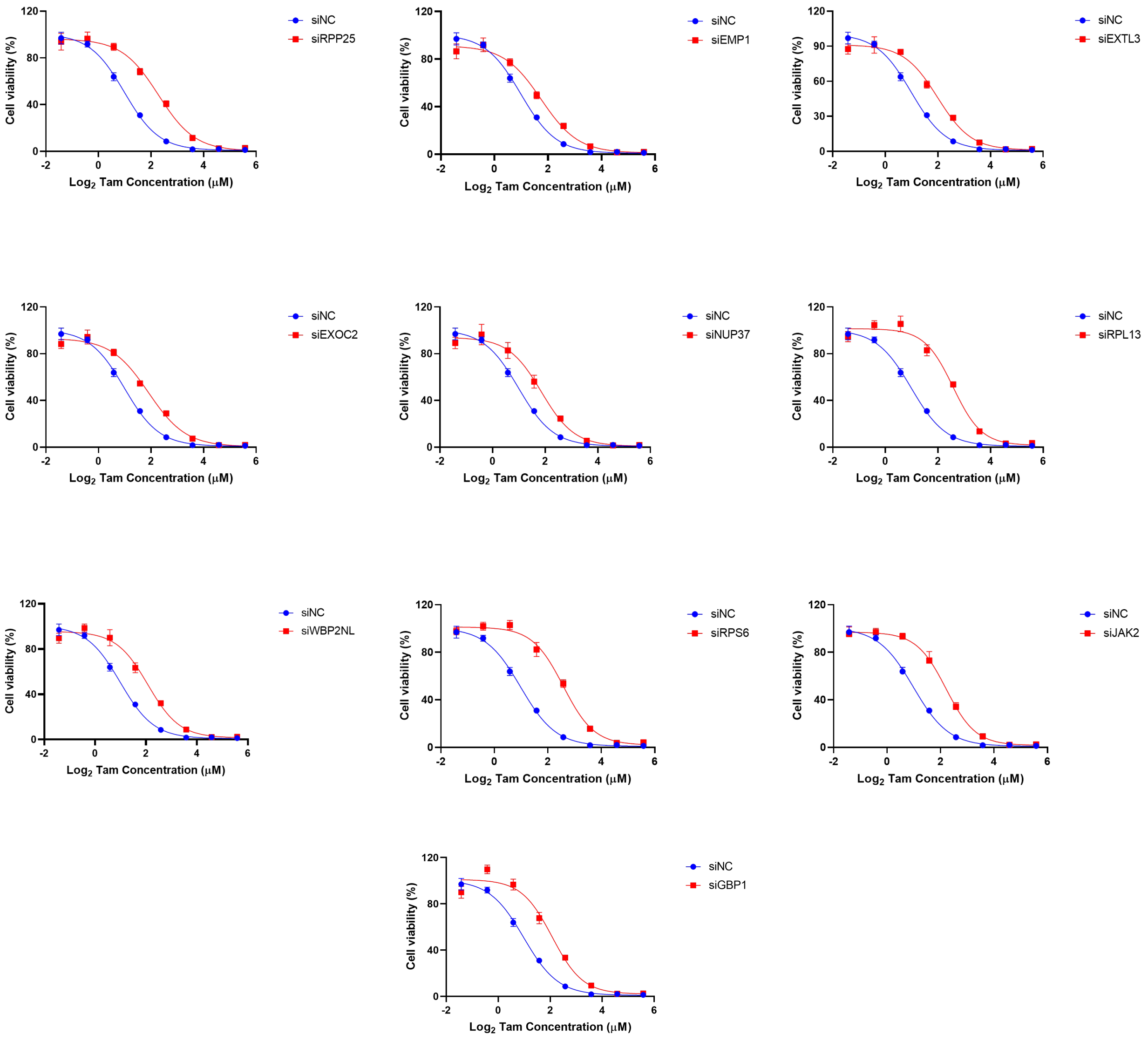


**Supplementary Figure S6**. Tamoxifen dose-response curves corresponding to the siRNA knockdown of 10 genes identified by TINDL in MCF7 cells. The p-values are calculated using an extra sum-of-squares F test.

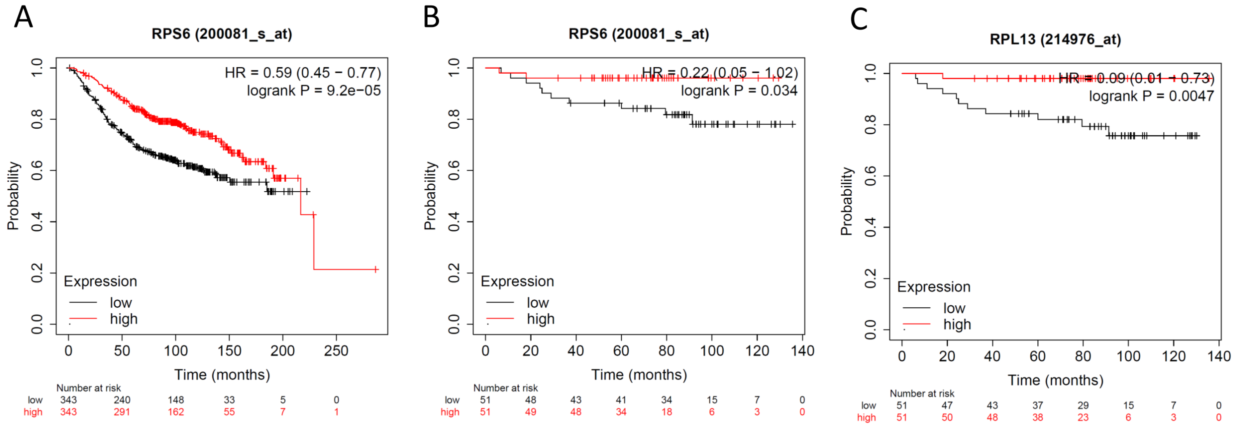


**Supplementary Figure S7**. Kaplan-Meier survival analysis of RPS6 and RPL13 gene expression in estrogen receptor positive/HER2 negative breast cancer patients using Kaplan-Meier plotter. A) Relapse free survival (RFS) of RPS6 in systemically untreated patients. B) Relapse free survival (RFS) of RPS6 in tamoxifen treated patients. C) Relapse free survival (RFS) of RPL13 in tamoxifen treated patients.
